# Coordinated orienting behavior emerges following unilateral optogenetic activation of Superior Colliculus GABAergic neurons

**DOI:** 10.64898/2026.10.27.761971

**Authors:** Hajime Suyama, Tatiana Lupashina, Myriam Hamon, Jérémie Sibille, Kai Lun Teh, Katharina Liebert, Matthew E. Larkum, Robert N. S. Sachdev, Jens Kremkow

## Abstract

Orienting coordinates movements of the eyes, whiskers, nose, and other effectors, but how the superior colliculus (SC) recruits these components over time remains unclear. We combined unilateral optogenetic activation of SC GABAergic neurons with high-speed tracking of facial movements and Neuropixels recordings to examine SC-evoked orienting in head-fixed mice. Stimulation recruited movements of the whiskers, nose, eyes, and pupil. Horizontal eye and nose movements were directed contralateral to the stimulated hemisphere, whereas vertical eye movements, pupil dilation, and whisker protraction did not differ significantly between stimulation sides. Nose and whisker movements generally preceded eye movements, followed by slower pupil dilation, revealing a temporally organized response. Recordings from the SC contralateral to the stimulated hemisphere revealed heterogeneous neuronal responses comprising short-latency excitation and delayed excitation or inhibition. Short-latency excitation was similar on trials with and without evoked movements, whereas delayed excitation was greater on trials that evoked eye or nose movements. Neurons modulated during spontaneous nose movements showed corresponding optogenetically evoked responses, linking stimulation-induced activity to endogenous movement-related activity. Thus, activation of SC GABAergic neurons initiates evolving population activity that recruits a coordinated sequence of orienting movements.

## Introduction

Animals orient toward salient events by coordinating movements of the eyes, head, nose, whiskers, and body. The superior colliculus (SC; optic tectum in non-mammalian vertebrates) is a conserved sensorimotor structure that transforms sensory information into spatially directed actions^1–7^. Classical stimulation and lesion studies established that the SC contains topographically organized representations linking locations in sensory space to orienting movements and attentional selection^8–14^. Importantly, SC stimulation can recruit movements extending beyond the eyes, including movements of the head, pinnae, whiskers, and body, indicating that collicular output can engage coordinated actions across multiple motor systems^15–17^.

Recent studies in mice have provided causal tests of how these actions are organized. Optogenetic stimulation revealed a three-dimensional representation of head-movement vectors within the SC, while genetically defined neuronal populations form functional modules associated with distinct forms of spatial orienting^18,19^. Cell- and projection-specific manipulations have further identified SC circuits that recruit approach, hunting, escape, and freezing, demonstrating that collicular output is organized according to both movement direction and behavioral function^20,21^. The activity of SC motor neurons is aligned with the direction of visual motion, with inhibitory gating regulating transmission through collicular premotor circuits^22^.

Recordings during natural behavior likewise show that SC activity accompanies eye, head, whisker, facial, and locomotor movements^23–25^. Rodent orienting nevertheless presents a distinct coordination problem. Unlike primates, mice make relatively small eye movements and rely heavily on movements of the whiskers, nose, head, and body to sample their surroundings. These effectors are coupled during spontaneous behavior: whisker positioning and attempted head movements are coordinated with movements of the eyes and face^26–28^. Recent work has shown that SC motor neurons are aligned with kinetic visual features and that inhibitory gating regulates transmission through collicular premotor circuits^22^. Yet it remains unclear how activation of an SC circuit recruits these effectors over time. In particular, it is not known whether the individual components are recruited simultaneously or sequentially, how their direction depends on the activated hemisphere, or which SC population responses distinguish trials with and without an overt orienting response.

Here, we combined unilateral optogenetic activation of SC GABAergic neurons with high-speed tracking of the eyes, pupil, nose, and whiskers and Neuropixels recordings from the contralateral SC. SC activation recruited a temporally organized sequence in which nose and whisker movements generally preceded eye movements. Horizontal eye and nose movements depended on the stimulation side, whereas other components were less strongly lateralized. Although stimulation produced rapid responses in the contralateral SC, delayed excitation was more strongly associated with trials in which eye or nose movements were evoked. Neurons modulated during spontaneous nose movements showed corresponding optogenetically evoked responses, linking stimulation-induced activity to endogenous movement-related activity. These findings show that focal SC activation initiates evolving population activity that recruits multiple components of an orienting action.

## Methods

### Animals and surgery

All procedures were performed in accordance with protocols for the care and use of laboratory animals approved by the Charité–Universitätsmedizin Berlin, Humboldt Universität-Berlin, and the Berlin Landesamt für Gesundheit und Soziales (LaGeSo). Adult transgenic VGAT-ChR2 mice of either sex (Slc32a1-COP4∗H134R/EYFP, The Jackson Laboratory, n = 7) were used. Animals were housed in groups of three and kept on a reverse 12-hour day-night cycle to ensure activity during behavioral training and adequate rest.

Mice were anesthetized with isoflurane (∼2 % in O_2_). Once mice were in the stereotaxic frame (Narishige, Japan), an eye ointment Bepanthen (Bayer, Germany) was applied and a local analgesic Lidocaine (100 μl) was injected under the skin. In the first step, lightweight aluminium headposts were affixed to the skull using Rely X (Applicap, 3M, US) followed by Paladur (Henry Schein, UK)^26^. Post-surgery analgesia was provided by a combination of Buprenorphine (0.1 mg/kg) and Carprofen (5 mg/kg), and in the days after surgery, analgesia was provided by Carprofen (5 mg/kg).

One week after surgery, mice were habituated to being handled and habituated to the behavioral apparatus. In the first days of habituation, mice were acclimated to having their head post handled by the experimenter and were acclimated to several minutes of head fixation. The duration of head fixation was gradually increased from 5 to 40 minutes. During head fixation habituation, mice also became accustomed to having their whiskers puffed, painted, or both. Once animals had been habituated, a second surgery, a craniotomy was made at AP 4.2 mm and ML ± 1.5 mm from bregma, aiming bilateral SC one day before the start of the experiment. A well was formed by layering Tetric Evoflow A1 (Ivoclar, Leichtenstein) around the craniotomy. The well was protected with silicon elastomer sealant Kwik-Cast (WPI, USA).

### Behavioral experiment setup

To enable precise optogenetic stimulation, electrophysiological recordings, and high speed behavior tracking, experiments were performed on the setup where a headpost, micromanipulators, and 4 cameras were placed using Thorlabs parts. A general layout (Air-Track) was adopted from previous studies^26,27,29^ in which a mouse is head-fixed on a floating platform with an air table. While mice are head-fixed, they still can exhibit naturalistic orienting behavior. Multi-Probe Micro-Manipulator System with two manipulators (New Scale Technologies, USA) was positioned behind the animal. Cameras (Basler ace U, Basler, Germany) were mounted on Fisso arms for precise positioning. Two monochrome cameras with the objectives with f = 50 mm and the IR filter (Navitar, USA) were aimed to eyes, one monochrome camera (f = 25 mm and the IR filter) was positioned in front of animals, and one color camera (f = 25 mm) was hung from the top to visualize whiskers and the platform movement. Eyes and the front view were illuminated by IR lights (Thorlabs, Germany). Whiskers were painted with UV paints and illuminated by an UV torch. Every frame acquisition was triggered by TTLs generated by a custom made Python script at 150 Hz.The TTLs were splitted and sent to an acquisition board (BNC-2110, National Instruments, USA) for synchronization. Obtained videos were recorded by PylonRecorder2 (https://gitlab.mpcdf.mpg.de/mpibr/scic/pylonrecorder/PylonRecorder2) and saved in an avi format.

### Optogenetic stimulation

The optic fiber (NA = 0.22, 200 µm core diameter, Thorlabs) was slowly lowered (5 μm/s) through the craniotomy aiming at the deep layer of the SC. Once approx. 2000 μm of insertion was achieved, LED pulses (470 nm, PlexBright, Plexon, USA) were applied and orofacial movements were simultaneously monitored by the experimenter (Pylon viewer, Basler). We adjusted the optic fiber placement to which obvious movements were evoked reliably. Stimulation intensity was ∼3 mW at the fiber tip and pulses were square continuous waveforms with duration of at least 1 s. Stimulation was triggered manually with intervals of at least ten seconds.

### Electrophysiology

Neural activity was recorded using high-density Neuropixels 1.0 probes (IMEC, Belgium) in the contralateral hemisphere of the optogenetic stimulation site. Data was amplified and digitized at 30k Hz in the Neuropixels probes and recorded through the PXIe system (NI-PXIe1071, National Instruments) and Open Ephys GUI (Open Ephys, https://open-ephys.org/gui). On the day of recording Kwik-Cast covering their craniotomy was carefully removed and the area was moistened by filling the recording-well with filtered phosphate buffered saline. A probe dyed with Di solution from Thermofisher Vybrant™ Multicolor Cell-Labeling Kit (Thermofisher, Germany) was positioned such that the tip of the probe just contacted the brain. The probe was advanced into the brain slowly at a speed of 5 μm/s, to a depth of 4500 μm and retracted for 50 μm. The probe was allowed to settle for ten minutes. At the end of each recording session, the probe was slowly (5 μm/s) retracted from the brain and the craniotomy was sealed with Kwik-Cast.

### Track Reconstruction

Mice were deeply anesthetized with isoflurane and transcardially perfused with 0.1% PBS and subsequently with 4 % paraformaldehyde (PFA) in 0.1 M phosphate buffer (PB, pH 7.2). Brains were carefully removed from the skull and post-fixed in 4 % PFA-PB overnight at 4 °C. Brains were embedded in 3 % agarose, for sectioning with a vibratome. All brains were sectioned at 100 µm. The sections were mounted on glass slides (Roti, Germany) using Mount FluorCare (Roti, Germany). They were imaged for track reconstruction using a microscope, Revolve (Echo by Bico, USA). Reconstruction was processed with a software HERBS (by Whitelock group, https://github.com/Whitlock-Group/HERBS). Channels in the putative SC were determined post-hoc by HERBS and distribution of baseline firing rates in multi-unit activity (MUA, see below).

### Orofacial movement tracking and analysis

Orofacial movements were tracked using python based DeepLabCut 3.0^30,31^. Pupils were tracked with eight points on the outline, nose positions were tracked with one point on the rhinarium, and whiskers were tracked with a point at the proximal end and another at the distal end of UV-illuminated parts of them on both sides of the head (**Figure 1B, Supplementary videos 1-3**). Tracked points were accepted at likelihood ≧ 0.9 for pupils and noses and ≧ 0.8 for whiskers. Pupil size and position as the center of mass were calculated when more than half of eight tracked points were available. Whisker angle was calculated when both proximal and distal points were available in the given frame.

**Figure 1.**
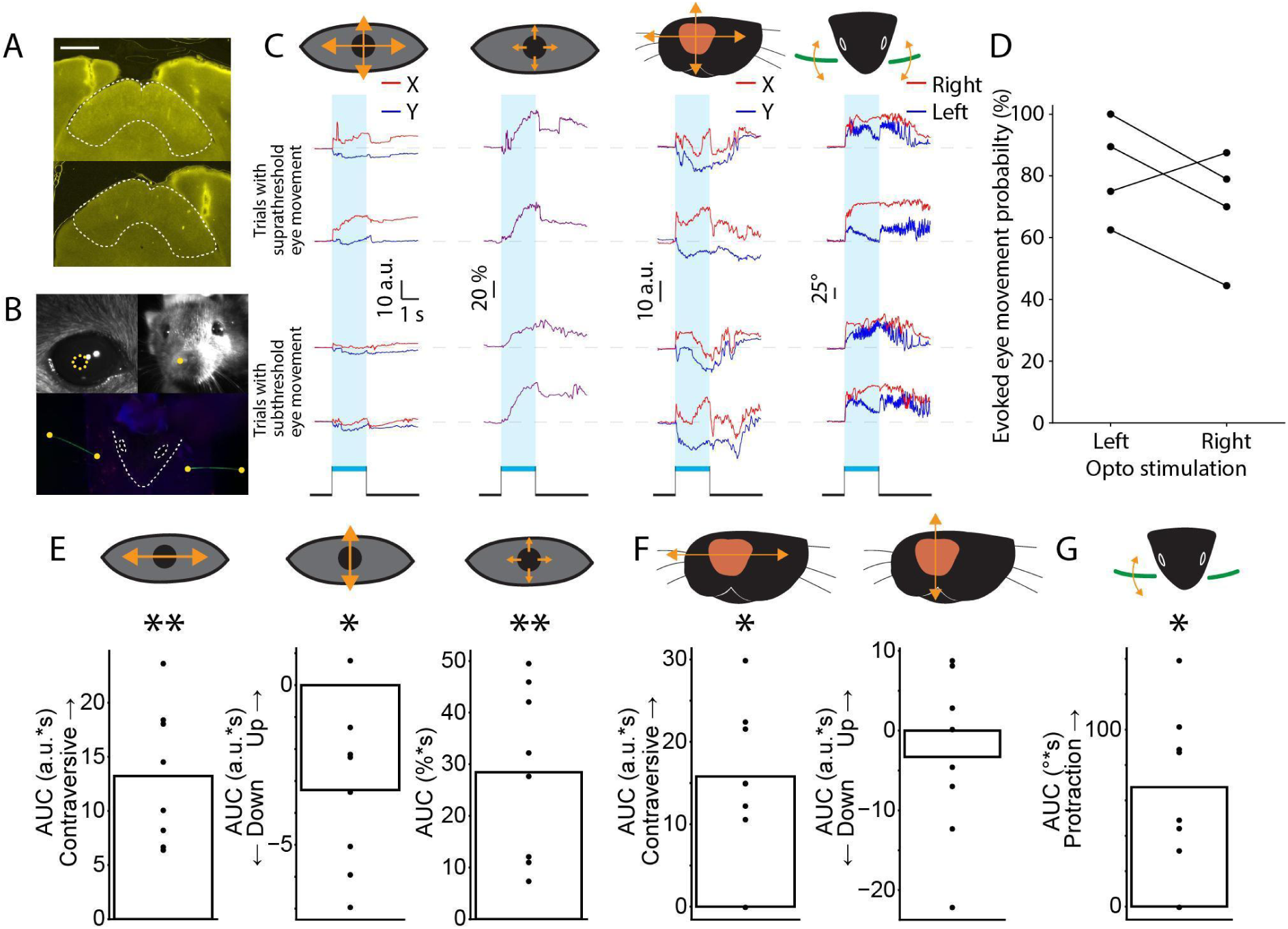
Unilateral optogenetic stimulation of the SC recruits multiple components of orienting behavior. **(A)** Representative histological sections from mice receiving stimulation of the left or right SC, showing ChR2 expression and optic fiber locations. Dashed lines delineate the SC. **(B)** Representative images illustrating behavioral tracking. Top left, eye image used to track horizontal and vertical eye position and pupil diameter; top right, facial image used to track nose position; bottom, UV-illuminated image used to track the painted left and right whiskers. Yellow dots indicate tracked landmarks used to train DeepLabCut. **(C)** Representative behavioral traces aligned to optogenetic stimulation. Blue shading and bars indicate the stimulation period (2 s). Trials were classified according to whether an eye movement was evoked (top) or not evoked (bottom). From left to right: horizontal (X, red) and vertical (Y, blue) eye position, pupil diameter, horizontal (X, red) and vertical (Y, blue) nose position, and right (red) and left (blue) whisker angle. Schematics indicate the corresponding movement axes. **(D)** Probability of evoking an eye movement following stimulation of the left or right SC. Connected points represent measurements from the same animal. **(E)** Stimulation-evoked changes in eye position and pupil diameter, quantified as the area under the curve (AUC) 0-2s. From left to right: horizontal eye displacement, with positive values indicating contraversive movement; vertical eye displacement; and change in pupil diameter. **(F)** Stimulation-evoked changes in nose position. Left, AUC of horizontal nose displacement, with positive values indicating contraversive movement; right, AUC of vertical nose displacement. **(G)** Stimulation-evoked whisker protraction, quantified as the AUC of the change in whisker angle. Bars indicate means across animals, and dots represent individual animals. Left- and right-SC stimulation probabilities in D were compared using a paired t-test. Values in E-G were compared with zero using Wilcoxon signed-rank tests. *p < 0.05, **p < 0.01.

Traces were cut from 1 s before the optogenetic stimulation until 10 s after that. Traces with less than 80 % of valid data points within ± 1 s of the optogenetic stimulation were discarded and the missing data points in valid trials were interpolated. X and Y coordinates of pupil and nose position were rotated to fit the axis of the inner-outer corners of the eyes and the axis of the right-left eyes, respectively. The value at 0 s was subtracted from traces followed by normalization (Δcoordinate). Then coordinates were normalized as: Normalized coordinate = Δcoordinate / distance of reference points * 100 where reference points for eye movements are the inner and outer corners of the eye and for nose movements they are inner corners of right and left eyes. Pupil size was normalized to the value at 0 s as changes in %. Quantified values were, onset (thresholds are 3 a.u. for eye and nose movement and 5 % and 5° for pupil size and whisker movement, respectively. At least two data points in a row should exceed the threshold), amplitudes, area under curve (AUC) during the stimulation, and peak time where the highest value during the stimulation was observed. Probabilities of evoked movement were calculated as the number of trials exceeding the threshold over the number of all valid trials in %. Sessions that have less than 5 trials wherein horizontal movements were evoked were rejected and only experiments that both hemisphere stimulation has enough eye-movement-evoked trials were included in comparison between stimulated hemispheres. Trials in which tracking of any orofacial part is failed and/or eye movement was not evoked were rejected for behavior sequence analysis. Trials in which tracking of any orofacial part is failed were rejected for investigating association of different orofacial part behaviors. Trials in which the given orofacial part movements were not evoked were rejected for investigating association between latencies and amplitudes.

For spontaneous orofacial movements, firstly invalid data points were interpolated throughout the session followed by rotation of X and Y coordinates of pupil and nose positions and normalization of their distance from the mean position to the eye size and inter-eye distance. Values were further normalized as (value - median value) / median value * 100 to have a controlled baseline value throughout the session. To detect fast eye movements, the threshold of the velocity was adopted from Bergmann et al.^27^, 0.1 a.u./frame. The same threshold was used for fast nose movement detection. 4°/frame was used for fast whisker movement detection. Detection was outside of optogenetic stimulation (from 1 s before the stimulation to 5 s after was rejected) and detected movements occurring within 250 ms of the last detected movement and exceeding the threshold only for one frame were rejected. Quantification of spontaneous whisker movements, traces ± 1 s of the fast movement onset were cut and the value at 0 s was subtracted from traces. AUC was quantified within a 1-s window after onsets.

Pupil data were averaged between right and left eyes, because evoked-responses of both pupils were similar (Supplementary video 1). Representative videos were cut around the optogenetic stimulation (-1 to 10s) with adjustment of brightness and contrast by a custom made python script using OpenCV package (https://docs.opencv.org/4.13.0/d5/de5/tutorial_py_setup_in_windows.html), and edited by Movavi (https://www.movavi.com) to add subtitles for notifying the stimulation.

### Multi-unit activity (MUA) extraction and spike sorting

MUA extraction was performed using our custom-built script (https://github.com/KremkowLab/tangential_recording)^32^. Briefly, after the bandpass filter (Butterworth filter order 2, 0.3-3 kHz), spikes were detected with a threshold of four standard deviations each channel.

Kilosort4^33^ was used for automatic detection and clustering of spikes followed by curation of good single units using quality metrics and labeling by Bombcell^34^ (https://github.com/Julie-Fabre/bombcell) and manual inspection with Phy2 (https://github.com/cortex-lab/phy).

### Firing rate (FR) analysis

Peristimulus time histograms (PSTHs) and Perievent time histograms (PETHs) were computed with a bin size of 10 ms. Averaged traces were z-scored for comparison between units.

For optogenetic stimulation evoked responses quantification windows were: baseline, -1 to 0 s; fast responses, 0-10 ms; delayed responses, 100-600 ms from stimulation. For spontaneous movement triggered activity quantification windows were: baseline, -1 s to -500 ms; after onsets, 0-500 ms from detected movement onsets. Mean FRs within the windows were used for statistical analysis. Raw FRs (Hz) in each trial were used to examine modulation in individual units.

Latencies of the first spike following optogenetic stimulation of units of which evoked fast responses are significantly higher than the baseline were quantified and median latencies of each unit were calculated. Before computing the median latency of fast excited units, outliers (2 out of 7 units) were detected by modified z-scores using median and median absolute deviation with a threshold of ± 3.5 and were excluded. Spike probabilities were calculated as the fraction of trials with spikes in a given time bin (2 ms).

Upon spontaneous movements, excited, inhibited, no change units were individually determined statistically and pooled in groups and compared between groups. Optogenetic stimulation evoked responses were pooled in groups determined by spontaneous movements and stimulation-evoked delayed responses were compared between groups.

### Statistics

Statistical tests were done by a python package, pingouin (https://pingouin-stats.org/) and a python module, statsmodels (https://www.statsmodels.org/devel/index.html). After normality of the data was tested, Friedman test and Linear Mixed Effects Model (LMM) was used to test the overall effects and association, Wilcoxon signed-rank test (Wilcoxon), Mann-Whitney U test (MWU), and paired t-test were used to test the individual group comparisons. Holm-Bonferroni correction was done, if needed. P values for association of behaviors were corrected using Benjamini-Hochberg FDR (FDR). LMM was used to account for animal identities as a dependent factor, and for testing behavior association optogenetic stimulation sides were also included. Significance was accepted at p < 0.05. Values shown in the text are mean ± standard error of mean (SEM).

## Results

### Optogenetic stimulation recruits multiple components of orienting behavior

To determine how SC activation recruits orienting behavior, we unilaterally stimulated ChR2-expressing GABAergic neurons in the posterolateral SC (AP 4.2 mm, ML ± 1.5 mm from bregma, DV ∼2 mm) while tracking eye position, pupil diameter, nose position, and whisker angle (**Figure 1A-C, Supplementary videos 1-3,** videos are from a single right SC stimulation trial). Behavioral responses varied across trials: some stimulation events evoked robust movements, whereas others produced little or no overt response (**Figure 1C**). Mean probabilities of evoked movement in the sessions were: eye, 84.5 ± 8.6 %; nose, 93.8 ± 6.2 %; whisker, 100 % (n = 9 sessions from 5 mice, 11-24 trials per session). We tested the efficiency of stimulation by eye movements. Representative trials with evoked horizontal eye movements showed pronounced horizontal and vertical eye displacements, whereas trials without evoked horizontal eye movements showed little change in eye position in both axes (**Figure 1C**). In representative evoked trials, eye movements were accompanied by pupil dilation and movements of the nose and whiskers, suggesting that stimulation could recruit multiple components of an orienting response.

Across animals, unilateral SC stimulation elicited eye movements on the majority of trials. The probability of eliciting an eye movement did not differ between stimulation of the left and right SC (**Figure 1D**; left SC: 81.7 ± 8.2 %; right SC: 70.2 ± 9.3 %; paired t-test, p = 0.247; n = 8 sessions in 4 mice). Stimulation produced significant contraversive horizontal eye movements **(Figure 1E**, left; AUC (area under curve): 13.2 ± 2.1 a.u.*s; Wilcoxon, p = 0.008), downward eye movements (**Figure 1E**, middle; AUC: -3.3 ± 0.9 a.u.*s; Wilcoxon, p = 0.016), and pupil dilation (**Figure 1E**, right; AUC: 28.5 ± 5.4 %*s; Wilcoxon, p = 0.008). Stimulation also produced significant contraversive nose displacement (**Figure 1F**, left; AUC: 15.8 ± 3.0 a.u.*s; Wilcoxon, p = 0.016) and whisker protraction (**Figure 1G**; AUC: 67.5 ± 14.8 degrees*s; Wilcoxon, p = 0.016), but did not produce a consistent vertical nose displacement (**Figure 1F**, right; AUC: -3.3 ± 3.5 a.u.*s; Wilcoxon, p = 0.641). Thus, unilateral activation of SC GABAergic neurons recruited eye, pupil, nose, and whisker responses, indicating that SC stimulation engages multiple components of orienting rather than producing an isolated movement of individual orofacial parts.

### The direction of horizontal orienting movements depends on the stimulated hemisphere

We next asked whether the movements recruited by SC stimulation depended on the stimulated hemisphere. Responses were therefore analyzed separately following left and right SC stimulation (**Figure 2**). Horizontal eye movements depended strongly on the stimulation side (**Figure 2A**). Left SC stimulation evoked rightward eye movements, whereas right SC stimulation evoked movements in the opposite direction. This reversal was evident across individual animals and resulted in a significant difference in horizontal eye displacement between left and right SC stimulation (**Figure 2A**; right stimulation, -10.4 ± 4.0 a.u.*s; left stimulation, 16.1 ± 4.6 a.u.*s; paired t-test, p = 0.002; n = 4 mice). In contrast, vertical eye displacement did not differ significantly between stimulation sides (**Figure 2B**; left SC: -3.1 ± 2.1 a.u.*s; right SC: -3.5 ± 1.7 a.u.*s; paired t-test, p = 0.90; n = 4 mice). Stimulation of either hemisphere also produced pupil dilation, with no significant difference between sides (**Figure 2C**; left SC: 34.9 ± 12.0 %*s; right SC: 22.0 ± 11.3 %*s; paired t-test, p = 0.08; n = 4 mice).

**Figure 2.**
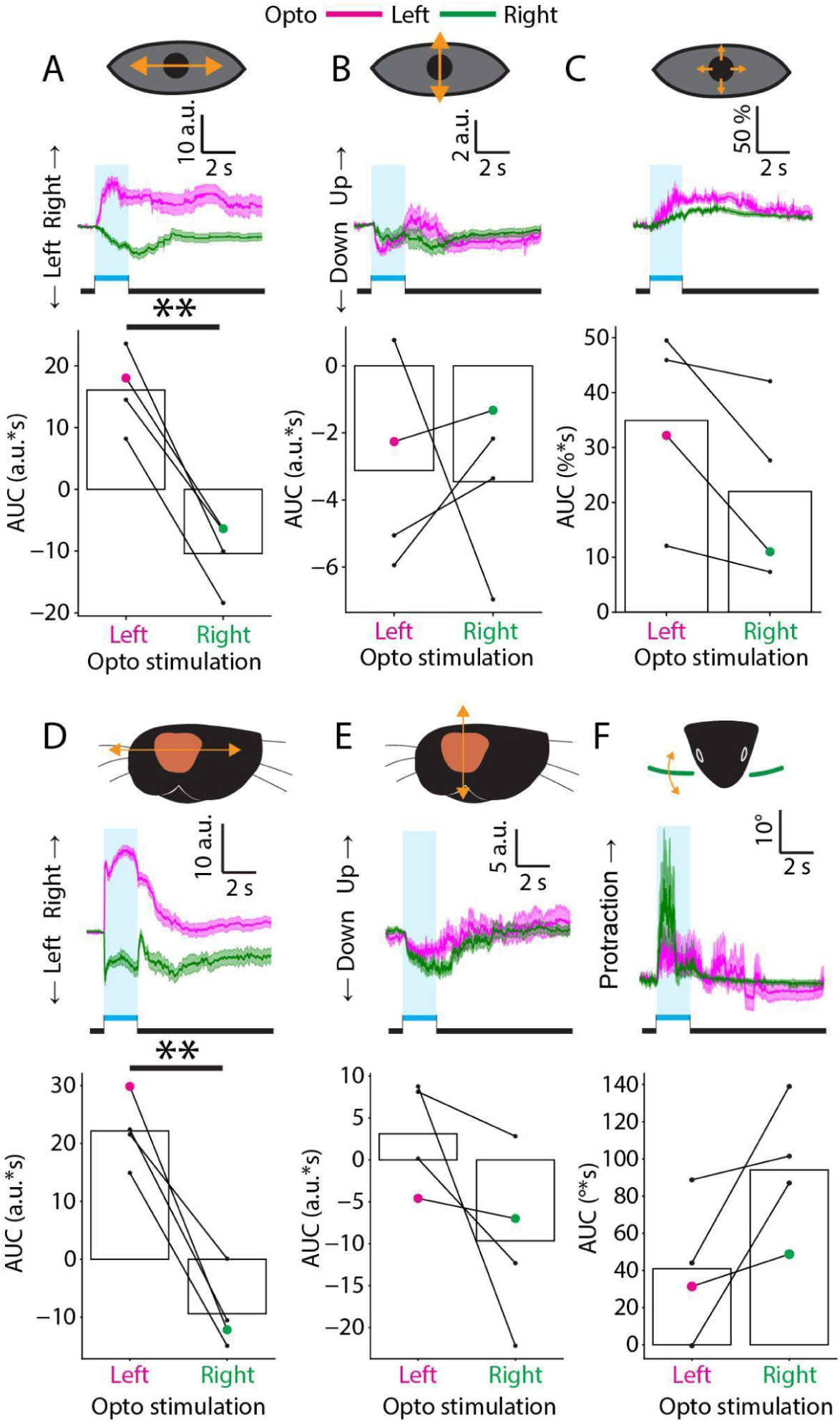
Stimulation side determines the direction of horizontal eye and nose movements. **(A)** Horizontal eye movements evoked by optogenetic stimulation of the left (magenta) or right (green) SC. Top, mean horizontal eye position aligned to stimulation onset. Bottom, area under the curve (AUC) for individual animals. Horizontal eye movements reversed direction with stimulation side, resulting in a significant difference between left and right SC stimulation. **(B)** Vertical eye movements following stimulation of the left or right SC. Top, mean vertical eye position aligned to stimulation onset. Bottom, AUC for individual animals. Vertical eye movements did not differ significantly between stimulation sides. **(C)** Changes in pupil diameter following stimulation of the left or right SC. Top, mean pupil diameter aligned to stimulation onset. Bottom, AUC for individual animals. Pupil dilation did not differ significantly between stimulation sides. **(D)** Horizontal nose movements following stimulation of the left or right SC. Top, mean horizontal nose position aligned to stimulation onset. Bottom, AUC for individual animals. Horizontal nose movements reversed direction with stimulation side, resulting in a significant difference between left and right SC stimulation. **(E)**Vertical nose movements following stimulation of the left or right SC. Top, mean vertical nose position aligned to stimulation onset. Bottom, AUC for individual animals.

Nose movements showed a similar separation between lateralized and non-lateralized components. Horizontal nose displacement reversed direction with stimulation side (**Figure 2D**), paralleling the horizontal eye movement and resulting in a significant difference between left and right SC stimulation (right stimulation, -9.4 ± 4.7 a.u.*s; left stimulation, 22.2 ± 4.3 a.u.*s; paired-t test, p = 0.005; n = 4 mice). In contrast, vertical nose movements did not differ significantly between stimulation sides (**Figure 2E**; right stimulation, -9.7 ± 7.4 a.u.*s; left stimulation, 3.1 ± 4.6 a.u.*s; paired t-test, p = 0.14; n = 4 mice).

Whisker protraction was evoked on both sides of the face following stimulation of either hemisphere. The magnitude of right whisker protraction did not differ significantly between stimulation sides (**Figure 2F**; left SC: 40.9 ± 26.2 degrees*s; right SC: 94.1 ± 26.4 degrees*s; paired t-test, p = 0.10; n = 4 mice). Similarly, left whisker protraction did not differ between stimulation sides (**Supplementary Figure 1A**; left SC: 45.0 ± 29.7 degrees*s; right SC: 49.3 ± 15.7 degrees*s; paired t-test, p = 0.90; n = 4 mice). Direct comparison of the two whiskers indicated that stimulation evoked approximately symmetric bilateral protraction (**Supplementary Figure 1B**), despite the asymmetry observed during spontaneous whisking (**Supplementary Figure 1C-F**), consistent with previous work^27^. Thus, the movements recruited by activation of SC GABAergic neurons differed in their relationship to the stimulated hemisphere. Horizontal eye and nose movements reversed direction with stimulation side, whereas vertical eye and nose movements, pupil dilation, and whisker protraction did not show significant stimulation-side dependence. SC stimulation therefore recruits multiple orienting components with distinct relationships to the stimulated hemisphere.

Vertical nose movements did not differ significantly between stimulation sides. **(F)** Right whisker angle following stimulation of the left or right SC. Top, mean whisker angle aligned to stimulation onset. Bottom, AUC for individual animals. Whisker protraction did not differ significantly between stimulation sides. Traces show mean ± SEM across trials from a representative recording session. Blue shading and blue bars indicate the optogenetic stimulation period. Black horizontal bars indicate the time window used to calculate the AUC. Bars in the lower plots indicate group means; dots and connecting lines represent paired measurements from individual animals. Colored dots identify the representative animal shown in the traces above. Left- and right-SC stimulation conditions were compared using two-sided paired t-tests (n = 4 mice). **p < 0.01.

### Optogenetic stimulation of the SC recruits temporally ordered sequences of orienting movements

SC stimulation recruited movements of the eyes, nose, and whiskers (**Figure 1**). We next asked whether these movement components were recruited simultaneously or in a temporal sequence. We therefore compared the onset latencies of horizontal eye movements, horizontal nose movements, and whisker protraction across 45 trials from 5 mice (**Figure 3A–C**). Nose movement was among the first detected components in 26 of 45 trials (58%; median latency from stimulation onset, 33.3 ms), and whisker protraction was among the first in 18 of 45 trials (40%; median latency, 46.7 ms). In contrast, eye movement was first in only 4 of 45 trials (9%; median latency, 233.3 ms). When two movements began within the same video frame, both were classified as occurring first; consequently, these percentages sum to more than 100%. Trial-by-trial onset latencies relative to eye-movement onset showed that nose movement and whisker protraction generally preceded eye movement, although the relative timing of nose and whisker movements varied across trials (**Figure 3A**). Analysis of the complete movement order identified three predominant sequences (**Figure 3B**). Whisker–nose–eye and nose–eye–whisker sequences each occurred in 14 of 45 trials (31%), whereas the nose–whisker–eye sequence occurred in 8 of 45 trials (18%). Together, these three sequences accounted for 80% of the analyzed trials. Representative traces illustrate each of these sequences (**Figure 3C**). Thus, SC stimulation did not recruit the three movement components simultaneously or in a single invariant sequence. Instead, nose and whisker movements typically preceded eye movements, while their relative order varied across trials.

**Figure 3.**
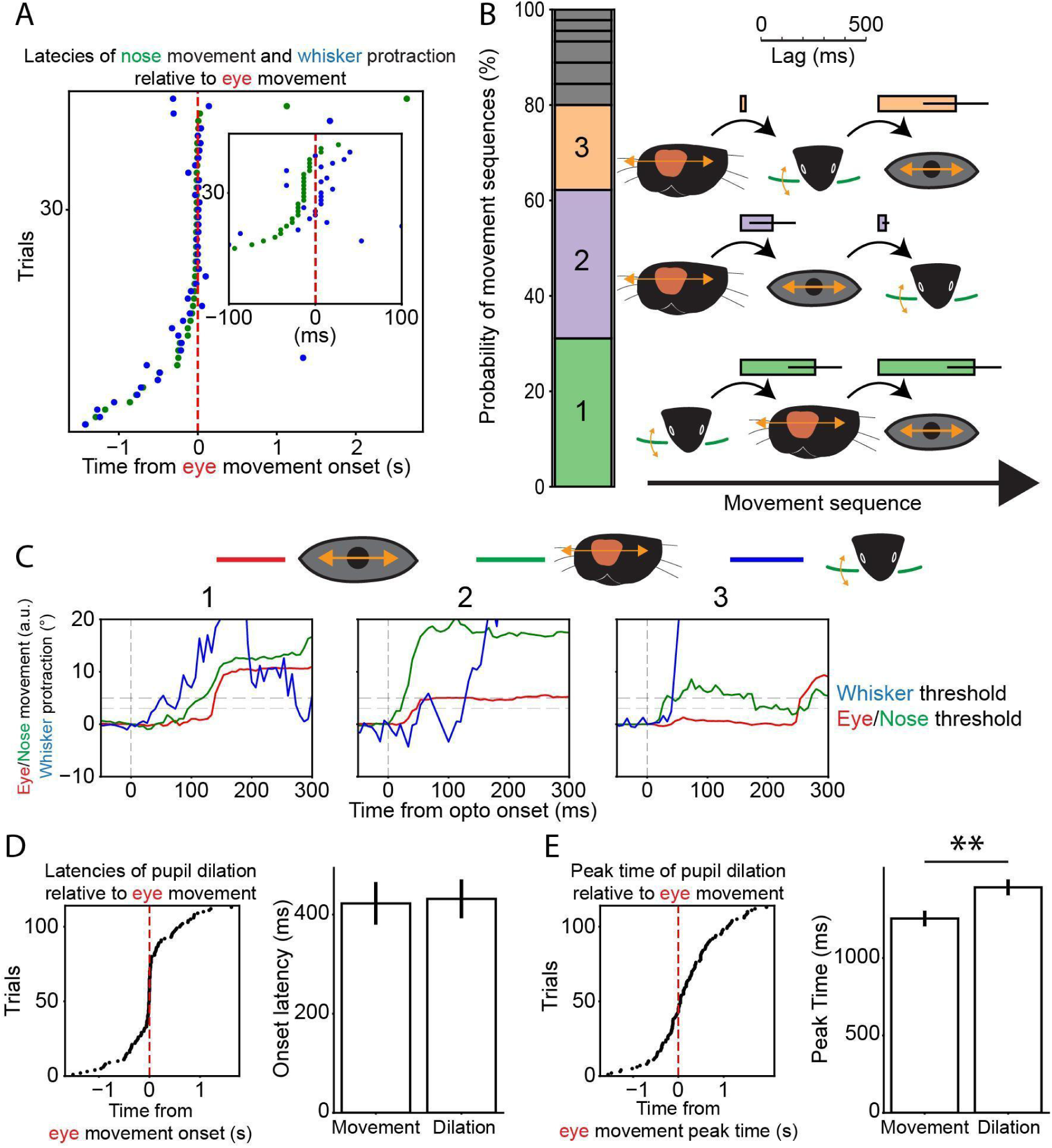
SC stimulation recruits a temporally ordered sequence of orienting movements. **(A)** Trial-by-trial onset latencies of nose movement (green) and whisker protraction (blue) relative to eye-movement onset. Each row represents one trial. The red dashed line indicates eye-movement onset; negative values indicate that nose or whisker movement preceded the eye movement. The inset shows the distribution of latencies around eye-movement onset at an expanded timescale. **(B)** Frequencies and temporal organization of movement sequences evoked by SC stimulation. The stacked bar shows the percentage of trials exhibiting each of the three most frequent movement sequences: whisker-nose-eye (green, sequence 1), nose-eye-whisker (purple, sequence 2), and nose-whisker-eye (orange, sequence 3). Gray segments indicate less frequent movement sequences. Schematics show the order of movements in the three principal sequences. Bars between successive movements show the mean latency difference ± SEM. Numbers identify the representative trials shown in C. **(C)** Representative traces illustrating the three principal movement sequences shown in A. Eye position is shown in red, nose position in green, and whisker angle in blue. Traces are aligned to optogenetic stimulation onset, indicated by the vertical dashed line. Horizontal dashed lines indicate the thresholds used to detect eye and nose movement onset and whisker protraction onset. Numbers correspond to the sequence categories shown in A. n = 45 trials from 5 mice. **(D)** Trial-by-trial onset latencies of pupil dilation relative to eye-movement onset (left). Mean ± SEM of onset latencies of eye movement and pupil dilation (right). **(E)** Trial-by-trial peak time of pupil dilation relative to eye-movement peak time (left). Mean ± SEM of peak time of eye movement and pupil dilation (right). Values from eye movements and pupil dilation were compared using LMM accounting for animal identities and stimulation sides. **p < 0.01. n = 113 trials from 5 mice.

Pupil dilation followed a somewhat different temporal profile. The onset latency of pupil dilation did not differ from that of horizontal eye movement (**Figure 3D**; pupil, 431.9 ± 39.2 ms; eye, 422.8 ± 43.0 ms; LMM, p = 0.858; n = 113 trials from 5 mice). However, pupil dilation reached its peak significantly later than horizontal eye movement (**Figure 3E**; pupil, 1457.1 ± 50.9 ms; eye, 1255.6 ± 50.5 ms; LMM, p = 0.005; n = 113 trials from 5 mice). Thus, although pupil dilation and horizontal eye movement were initiated at similar times, pupil dilation evolved more slowly and reached its maximum later.

### SC-evoked movement components show partial trial-by-trial covariation

Having established that SC stimulation recruits temporally ordered movements (**Figure 3**), we next asked whether the timing and amplitude of the individual movement components covaried across trials. For horizontal eye movements and horizontal nose movements, but not for whisker protraction, earlier movement onset was associated with greater movement amplitude (**Figure 4A, B, C**; horizontal eye movement: LMM with FDR, slope = -7.71, p < 0.001, n = 122 trials from 5 mice; horizontal nose movement: LMM with FDR, slope = -4.95, p = 0.018, n = 134 trials from 5 mice; whisker protraction: LMM with FDR, slope = -10.53, p = 0.284, n = 55 trials from 5 mice; Trials in which movement of the given facial part was evoked). Thus, trials with shorter response latencies tended to produce larger movements. However, this relationship was not found for whisker protraction. This could be because whisker protraction has less variety in its onset latencies (**Figure 4C**).

**Figure 4.**
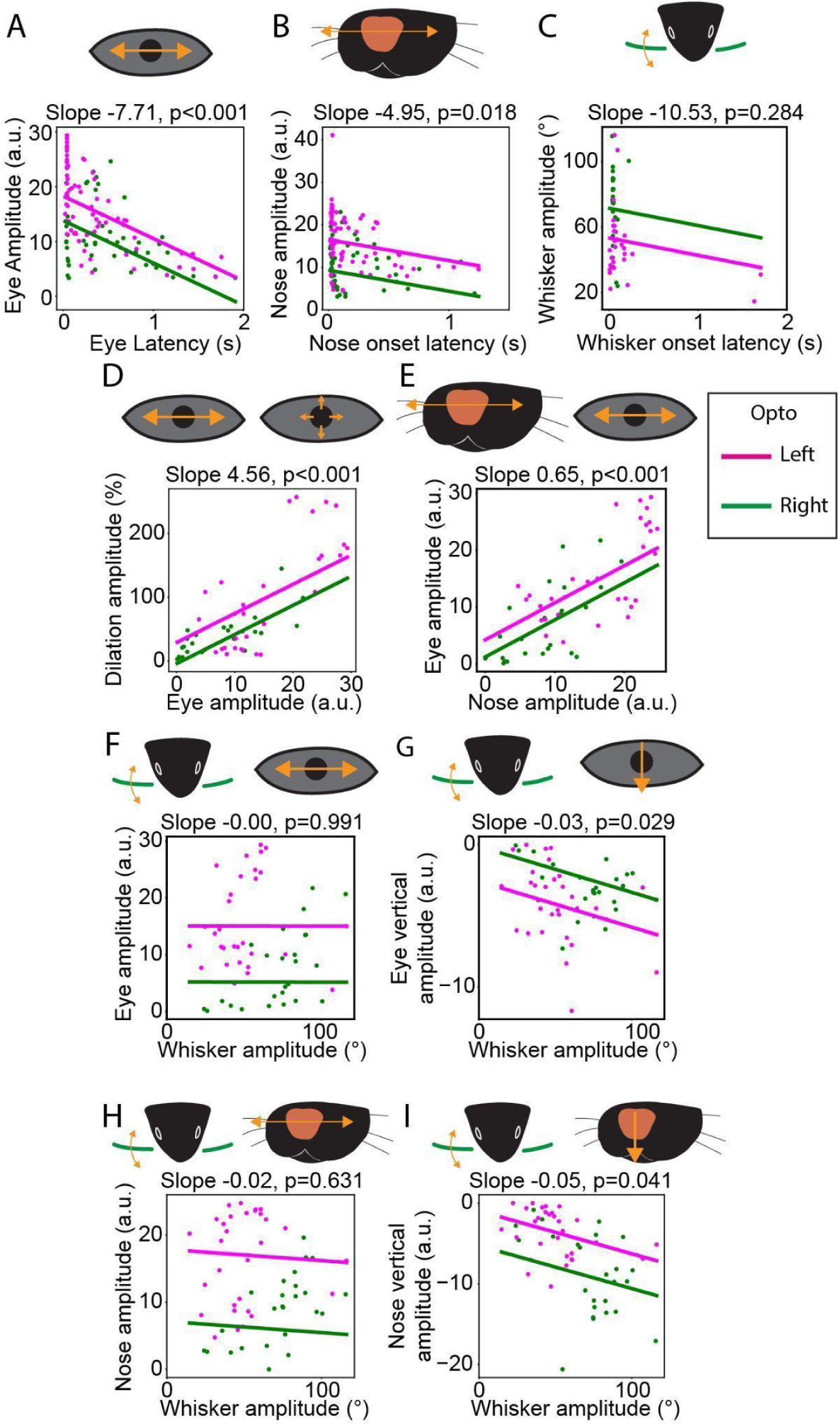
Relationships between the timing and amplitude of SC-evoked movement components. **(A)** Relationship between horizontal eye-movement onset latency and horizontal eye-movement amplitude. **(B)** Relationship between horizontal nose-movement onset latency and horizontal nose-movement amplitude. **(C)** Relationship between whisker-protraction onset latency and whisker-protraction amplitude. **(D)** Relationship between horizontal eye-movement amplitude and pupil-dilation amplitude. **(E)** Relationship between horizontal nose-movement amplitude and horizontal eye-movement amplitude. **(F)** Relationship between whisker-protraction amplitude and horizontal eye-movement amplitude. **(G)** Relationship between whisker-protraction amplitude and downward eye displacement. **(H)** Relationship between whisker-protraction amplitude and horizontal nose-movement amplitude. **(I)** Relationship between whisker-protraction amplitude and downward nose displacement. Horizontal eye and nose amplitudes represent contraversive displacement. Negative vertical eye and nose values indicate downward movement. Whisker amplitudes represent protraction. Points represent individual trials (n = 122 trials (A), n = 134 trials (B), n = 55 trials (C-I), 5 mice). Solid lines show LMM regression fits and colors represent stimulation sides (magenta, left hemisphere; green, right hemisphere). Associations were tested by LMM regression accounting for animal identities and stimulation sides. LMM regression slope and corresponding p values are shown above each panel. P values were corrected for 9 comparisons using Benjamini-Hochberg FDR.

The amplitudes of several simultaneously recruited components also covaried. Trials with valid tracking in all facial parts were used in the comparisons, regardless if movements exceeded the thresholds. Horizontal eye-movement amplitude was positively associated with pupil dilation (**Figure 4D**; LMM with FDR, slope = 4.56, p < 0.001, n = 55 trials from 5 mice) and with horizontal nose-movement amplitude (**Figure 4E**; LMM with FDR, slope = 0.65, p < 0.001, n = 55 trials from 5 mice). In contrast, whisker-protraction amplitude was not associated with either horizontal eye-movement amplitude (**Figure 4F**; LMM with FDR, slope = -0.00, p = 0.991, n = 55 trials from 5 mice) or horizontal nose-movement amplitude (**Figure 4H**; LMM with FDR, slope = -0.02, p = 0.631, n = 55 trials from 5 mice). Whisker-protraction amplitude was associated with eye downward or nose downward displacement, with stronger protraction associated with greater downward eye (**Figure 4G**; LMM with FDR, slope = -0.03, p = 0.029, n = 55 trials from 5 mice) or nose movement (**Figure 4I**; LMM with FDR, slope = -0.05, p = 0.041, n = 55 trials from 5 mice).

### Optogenetic stimulation evokes rapid and delayed neural responses in the contralateral SC

Computational analysis indicated that stimulation of SC inhibitory neurons is likely to lead to contraversive choices, if intercollicular inhibition overpowers the local inhibition^35^. We then next examined the neural activity in the SC contralateral to the stimulation site associated with optogenetically evoked orienting behavior using Neuropixels (**Figure 5A, Supplementary figure 2**). Multi-unit activity revealed a rapid increase in firing immediately following stimulation, followed by a broader delayed response lasting several hundred milliseconds (**Figure 5B**, **Supplementary figure 3A and B**). Similar response structure was observed at the single-unit level, where optogenetic stimulation recruited heterogeneous patterns of excitation and suppression across neurons (**Figure 5C**). A subset of neurons exhibited excitation within 10 ms of stimulation onset. First-spike latencies in these neurons were tightly locked to stimulation, with a median latency of 8.1 ms across the population and 3.0 ms in the example neuron (**Figure 5D**). These short-latency responses demonstrate rapid recruitment of the contralateral SC following unilateral stimulation, although the underlying synaptic pathway needs to be investigated in future.

**Figure 5.**
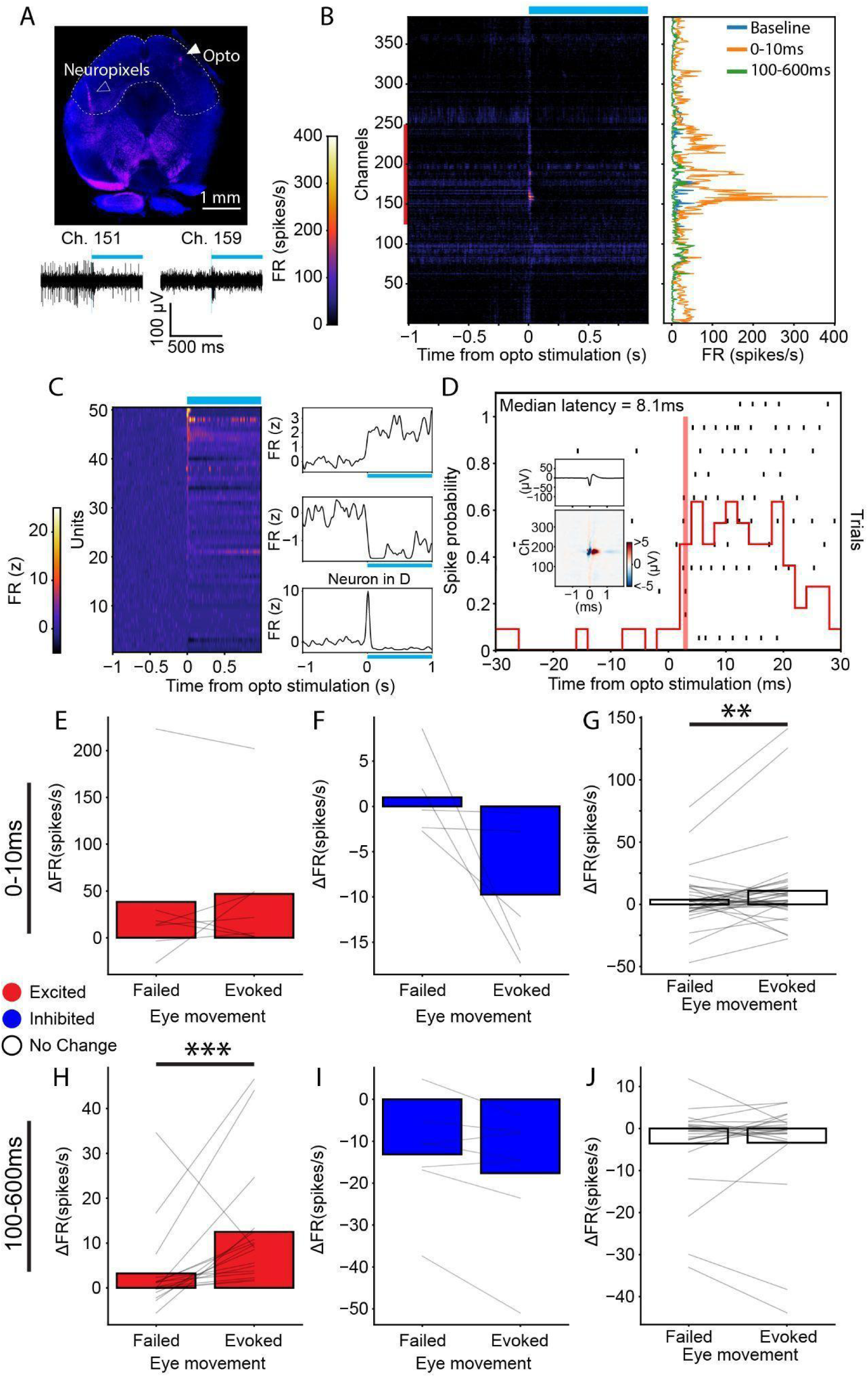
Neuropixels recordings reveal rapid and delayed responses in the SC contralateral to stimulation. **(A)** Top, representative histological section showing the Neuropixels probe track in the recorded SC and the optogenetic stimulation site in the contralateral SC. DAPI is shown in blue, and insertion tracks are shown in magenta. Bottom, representative band-pass-filtered voltage traces recorded across Neuropixels channels during optogenetic stimulation. Blue lines indicate stimulation pulses. **(B)** Multi-unit activity aligned to stimulation onset. Left, firing-rate heat map across probe channels, ordered by recording depth. The red line indicates channels located within the SC. Heat map represents firing rates ranging 0 to 400 Hz. Right, mean multi-unit firing rates during baseline window from -1 to 0 s from stimulation onset, the early response window from 0 to 10 ms, and the delayed response window from 100 to 600 ms after stimulation onset. **(C)** Single-unit activity aligned to stimulation onset. Left, each row represents one neuron, and color indicates z-scored firing rates ranging -5 to 25 z. Neurons are ordered by z-scored firing rates in the 0-to-10 ms bin. Stimulation recruited heterogeneous responses comprising short-latency excitation and delayed excitation or inhibition; right, representative smoothed PSTHs. Top to bottom, an excited unit, an inhibited unit, a unit with fast excitation which is further shown in panel D. **(D)** Representative neuron with short-latency excitation. Top, spike raster aligned to stimulation onset; bottom, spike probability calculated with a binsize of 2 ms. The red vertical line indicates the median first-spike latency of the example neuron, 3 ms. Across neurons classified as excited during the 0 to 10 ms window, the median first-spike latency was approximately 8 ms (n = 5 neurons). Insets show the mean waveform on the channel with the largest waveform amplitude and the waveform across neighboring channels. The heat map represents potentials ranging < 5 to > 5 µV. **(E-G)** Bar plot represents mean firing rate changes in the 0-to-10 ms bin from baseline in trials in which eye movement was not evoked and was evoked of units that were significantly excited (E), were significantly inhibited (F), and were not modulated (G). Line plots represent individual unit data. **(H-J)** Bar plot represents mean firing rate changes in the 100-to-600 ms bin from baseline in trials in which eye movement was not evoked and was evoked of units that were significantly excited (H), were significantly inhibited (I), and were not modulated (J). Line plots represent individual unit data. Data were obtained from 3 recording sessions in 3 mice. Comparisons between conditions were tested with LMM accounting for animal identities. **p < 0.01, ***p < 0.001.

Across three mice, most neurons were not significantly modulated relative to baseline during the 0–10 ms window, whereas seven neurons were excited and five were inhibited (**Supplementary Figure 3C**). Short-latency excitation did not differ between trials with and without evoked eye movements (**Figure 5E**; evoked, 47.0 ± 25.0 Hz; failed, 38.2 ± 29.2 Hz; LMM, p = 0.54). Short-latency inhibition also did not differ significantly between trial types (**Figure 5F**; evoked, −9.7 ± 3.0 Hz; failed, 1.0 ± 1.8 Hz; LMM, p = 0.08). Neurons classified as unmodulated relative to baseline nevertheless showed a small trial-type difference (**Figure 5G**; evoked, 10.9 ± 5.2 Hz; failed, 3.8 ± 3.3 Hz; LMM, p = 0.006). During the delayed 100–600 ms window, 17 neurons were excited, seven were inhibited, and 26 were not significantly modulated relative to baseline. Delayed excitation was greater on trials with evoked eye movements than on failed trials (**Figure 5H**; evoked, 12.5 ± 3.2 Hz; failed, 3.2 ± 2.2 Hz; LMM, p < 0.001). Delayed inhibition and activity in neurons classified as unmodulated did not differ significantly between trial types (**Figure 5I,J**).

Similarly, short-latency excitation and inhibition did not differ between trials with and without evoked nose or whisker movements, as well as responses in unchanged neurons (nose excited: evoked, 50.4 ± 26.0 Hz; failed, 37.6 ± 29.5 Hz; LMM, p = 0.23; nose inhibited: evoked, −4.8 ± 2.7 Hz; failed, 2.0 ± 2.8 Hz; LMM, p = 0.16; nose unchanged: evoked, 7.8 ± 3.4 Hz; failed, 7.1 ± 4.2 Hz; LMM, p = 0.44; whisker excited: evoked, 50.4 ± 26.7 Hz; failed, 27.1 ± 19.1 Hz; LMM, p = 0.53; whisker inhibited: evoked, −1.8 ± 0.8 Hz; failed, 7.2 ± 8.3 Hz; LMM, p = 0.08; whisker unchanged: evoked, 8.9 ± 3.6 Hz; failed, 10.9 ± 3.7 Hz; LMM, p = 0.81). Moreover, delayed excitation was greater on trials with evoked nose movements than on failed trials (evoked, 13.1 ± 3.1 Hz; failed, 3.0 ± 2.0 Hz; LMM, p < 0.001). The rest of the responses did not differ significantly (nose inhibited: evoked, −15.4 ± 4.1 Hz; failed, -14.1 ± 4.4 Hz; LMM, p = 0.83; nose unchanged: evoked, -3.6 ± 1.7 Hz; failed, -3.7 ± 2.0 Hz; LMM, p = 0.053; whisker excited: evoked, 11.3 ± 2.6 Hz; failed, 8.4 ± 2.5 Hz; LMM, p = 0.31; whisker inhibited: evoked, −15.0 ± 4.2 Hz; failed, -14.2 ± 4.1 Hz; LMM, p = 0.53; whisker unchanged: evoked, -3.7 ± 1.8 Hz; failed, -4.5 ± 2.1 Hz; LMM, p = 0.29).

Note that although responses to optogenetics stimulation varied, we didn’t find any significant differences in electrophysiological features between neurons showing different responses, except spike amplitudes where excited neurons had significantly larger spike amplitudes than inhibited or unchanged neurons in delayed responses (**Supplementary figure 3D-M**). The result is similar to the previous study^36^ where authors showed that no difference was found between opto-tagged GABAergic neurons and the rest in the superficial (visual) SC. Thus, unilateral activation of SC GABAergic neurons produced both rapid and delayed changes in contralateral SC activity. Eye-movement recruitment was most clearly associated with greater delayed excitation rather than with the initial excitatory response.

### Spontaneous movement-related activity is associated with optogenetically evoked SC responses

Because delayed stimulation-evoked activity varied with eye-movement recruitment, we asked whether optogenetic stimulation engaged neurons that also participated in spontaneous orofacial movements. For each spontaneous movement, neurons were classified as excited, inhibited, or unchanged relative to baseline. During spontaneous nose movements, 27 neurons were excited, 10 were inhibited, and 13 were unchanged. During spontaneous eye movements, 15 neurons were excited, 4 were inhibited, and 31 were unchanged. During spontaneous whisker protraction, 32 neurons were excited, 11 were inhibited, and 7 were unchanged (**Figure 6A,C,E**).

**Figure 6.**
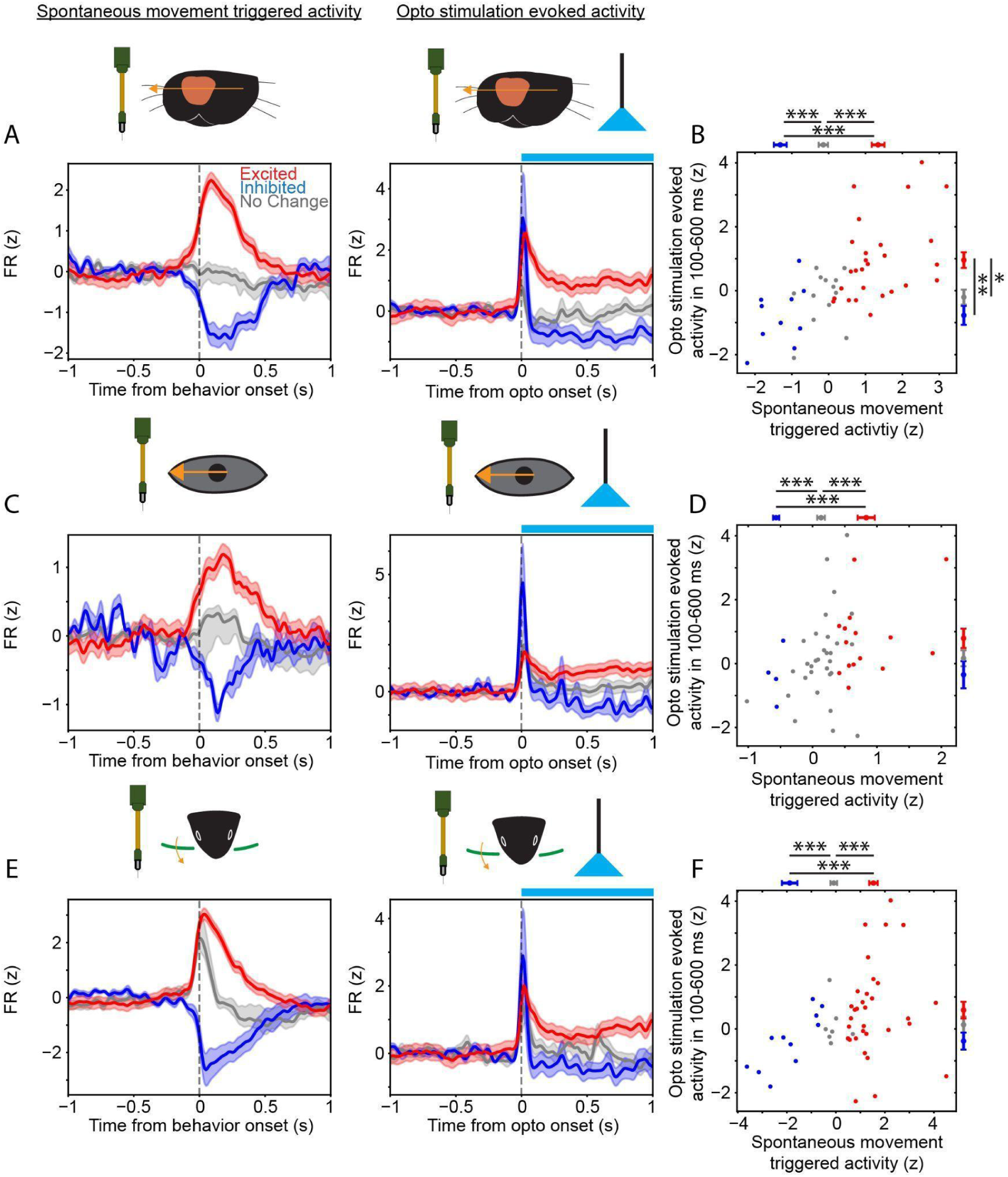
Spontaneous movement-related activity is associated with optogenetically evoked responses in the SC. **(A)** Left, mean firing-rate responses aligned to spontaneous nose-movement onset for neurons classified as excited, inhibited, or unchanged during spontaneous nose movement. Right, mean responses of the same neurons aligned to optogenetic stimulation onset. **(B)** Relationship between spontaneous nose-movement-related activity (X axis) and optogenetically evoked activity (Y axis). Each point represents one neuron. Marginal summaries show mean ± SEM for neurons classified as excited, inhibited, or unchanged during spontaneous nose movement. **(C)** Mean firing-rate responses aligned to spontaneous eye-movement onset and optogenetic stimulation onset for neurons classified according to their spontaneous eye-movement-related activity. **(D)** Relationship between spontaneous eye-movement-related activity (X axis) and optogenetically evoked activity (Y axis). **(E)** Mean firing-rate responses aligned to spontaneous whisker-protraction onset and optogenetic stimulation onset for neurons classified according to their spontaneous whisker-related activity. **(F)** Relationship between spontaneous whisker-protraction-related activity (x axis) and optogenetically evoked activity (Y axis). Red indicates neurons significantly excited during the corresponding spontaneous movement, blue indicates neurons significantly inhibited, and gray indicates neurons without a significant response relative to baseline. Traces show smoothed (Gaussian filter, sigma = 20 ms) mean ± SEM. Blue bars on top of traces upon optogenetic stimulation indicate the optogenetic stimulation period. Spontaneous movement-related activity was quantified in the 0 to 500 ms window, and optogenetically evoked activity was quantified in the delayed 100 to 600 ms window. Neurons (n = 50 from 3 mice) were classified individually using Friedman test followed by Wilcoxon signed-rank test with Holm-Bonferroni. Group comparisons were performed using LMM accounting for animal identities followed by MWU with Holm-Bonferroni. *p < 0.05, **p < 0.01, ***p < 0.001.

Across neurons, spontaneous movement-related activity was positively associated with activity during the delayed 100–600 ms stimulation window (**Figure 6B,D,F**; nose movement: LMM slope = 0.593, p < 0.001; eye movement: slope = 0.926, p = 0.001; whisker protraction: slope = 0.343, p = 0.001; n = 50 neurons from 3 mice). Thus, neurons that were more strongly modulated during spontaneous movements also tended to show greater modulation following optogenetic stimulation.

This relationship was most clearly expressed when neurons were grouped according to their spontaneous nose-movement responses. Optogenetically evoked activity differed among nose-excited, nose-inhibited, and nose-unchanged neurons (**Figure 6B**; omnibus LMM, p < 0.001). Nose-excited neurons showed greater stimulation-evoked activity than nose-inhibited neurons (p = 0.001) and nose-unchanged neurons (p = 0.03), whereas nose-inhibited and nose-unchanged neurons did not differ significantly (p = 0.10). Optogenetically evoked activity did not differ significantly among groups defined by spontaneous eye-movement responses (**Figure 6D**; LMM, p = 0.10). Optogenetically evoked activity differed overall among neurons classified by their spontaneous whisker responses (**Figure 6F**; LMM, p = 0.04). But note that none of the Holm-corrected pairwise comparisons was significant. These results link stimulation-evoked contralateral SC activity to endogenous movement-related activity, with the clearest group separation observed for neurons classified by their responses during spontaneous nose movements.

## Discussion

Here, we show that unilateral activation of GABAergic neurons in the mouse SC recruits a coordinated but variable orienting response involving the whiskers, nose, eyes, and pupil. The stimulated hemisphere determined the direction of horizontal eye and nose movements, whereas vertical eye and nose movements, pupil dilation, and whisker protraction did not differ significantly between stimulation sides. These components were not recruited simultaneously: nose and whisker movements generally began before eye movements, followed by slower pupil dilation, although the precise order varied across trials. Recordings from the contralateral SC revealed rapid and delayed neuronal responses. Short-latency excitation was similar on trials with and without evoked eye movements, whereas delayed excitation was greater on trials in which stimulation evoked an eye movement. Finally, neurons modulated during spontaneous orofacial movements showed corresponding optogenetically evoked responses, with the clearest group separation observed for spontaneous nose-movement-related activity. Together, these findings show that SC stimulation does not produce an isolated motor output but initiates an evolving network response that recruits multiple components of an orienting action.

### SC stimulation recruits multiple components of orienting behavior

These findings extend the classical view of the SC as a structure that transforms spatial information into orienting movements. Electrical stimulation and lesion studies established the central role of the SC in movements of the eyes, head, pinnae, and body and showed that collicular output can coordinate oculomotor, cephalomotor, and skeletomotor systems^14–17,37^. Much of the mechanistic literature, particularly in primates, has nevertheless focused on the relationship between SC activity and saccadic eye movements. Orienting, however, is an action of different motor systems rather than an eye movement alone. Our findings make this distinction explicit in mice: activation at one SC site recruited movements across several facial effectors, with the eye movement forming one component of a broader behavioral response.

Recent optogenetic studies in mice have revealed complementary principles of SC motor organization. Wilson et al.^18^ demonstrated a three-dimensional representation of head-movement vectors, whereas Masullo et al.^19^ identified genetically defined modules associated with distinct forms of spatial orienting. Cell- and projection-specific studies have further identified pathways that promote approach, hunting, escape, and freezing^20,21^. Together, these studies show that SC output is organized both spatially and according to behavioral function. The present study adds temporal and multi-effector dimensions to this organization. Rather than asking which movement vector or behavioral category is produced, we examined how a focal perturbation was expressed across several simultaneously monitored effectors. The resulting components differed in their relationships to the stimulated hemisphere, their onset times, and their trial-by-trial covariation. An SC signal may therefore not be copied identically to each motor system. Instead, collicular activation appears to initiate a process in which distinct outputs are recruited according to their own anatomical and temporal organization.

The distinction between strongly and weakly lateralized components was particularly informative. Horizontal eye and nose movements reversed with stimulation side, consistent with a directional signal linked to the spatial organization of the SC. By contrast, vertical eye and nose movements, pupil dilation, and bilateral whisker protraction did not differ significantly between stimulation sides. An orienting response may therefore combine a lateralized component specifying horizontal direction with less lateralized components related to facial positioning, sensory sampling, or arousal. The direction of the vertical component may instead depend on the position stimulated within the SC motor map: vertical head-movement vectors vary along the anterior-posterior and medial-lateral axes of the mouse SC^19^. Because the stimulation sites in the present study were relatively lateral and posterior, their location may have contributed to the consistent vertical response observed across hemispheres. However, because the number of animals and stimulation sites was limited, the absence of a significant side difference should not be interpreted as evidence of complete bilateral symmetry or as a definitive mapping of the vertical component.

### SC output pathways provide routes for coordinated orofacial movements

The anatomical organization of SC output provides plausible routes through which a focal perturbation could recruit these different components. Neurons in the intermediate and deep SC give rise to descending tectobulbar and tectoreticular projections that reach premotor regions throughout the midbrain, pons, and medulla^16,38,39^. GABAergic neurons in the intermediate SC include local and interlaminar interneurons as well as commissural and tectofugal neurons, indicating that subsets of these neurons contribute to long-range as well as local inhibition^40^. Classical gaze-control circuitry provides strongly lateralized routes for horizontal eye movements, whereas vertical eye movements depend more heavily on bilateral midbrain premotor networks. Although this organization has been established most extensively in primates, it provides a possible substrate for the stronger lateralization of horizontal than vertical eye displacement observed here. Moreover, different targets of tectoreticular projections control eye movement or head displacement, i.e., projections to the paramedian pontine reticular formation or the gigantocellular nucleus, respectively. The authors showed that the tectoreticular projections to two regions are collateralized^39^. Therefore, the same SC neurons can evoke eye and head movement. The SC also has direct access to orofacial motor circuits. Descending SC axons terminate in the facial nucleus and in medullary reticular regions containing facial premotor and rhythm-generating neurons, including the intermediate reticular formation and pre-Botzinger complex^41^. Commissural coupling within respiratory-whisking networks could allow a predominantly crossed output from one SC to recruit whisker movements on both sides of the face^41,42^. These projections also provide plausible routes for the coordinated nose and whisker movements observed here, although the specific premotor pathway controlling nose displacement has not been established. Pupil dilation probably reflects recruitment of a broader autonomic or arousal component of orienting rather than the lateralized somatic pathways controlling horizontal movements. Microstimulation of the deep optic tectum evokes pupil dilation in barn owls, and stimulation of the intermediate SC in monkeys can evoke pupil dilation at currents below the threshold for eliciting saccades^2,43^. Thus, SC activation can recruit the pupillary component of orienting without an overt eye movement. However, the downstream pathways through which SC activity engages sympathetic and parasympathetic pupil-control circuits remain incompletely defined^44^.

Recent work suggests that SC-dependent orienting also contributes to social behavior. SC neurons projecting to the ipsilateral ventral tegmental area (VTA) are recruited during orienting toward unfamiliar conspecifics, and manipulation of this pathway disrupts social orienting and alters subsequent social interaction^4^. This pathway is also affected in models and clinical measures of autism spectrum disorder. Shank3 knockout mice show reduced activity and altered interneuronal coordination among SC-VTA-projecting neurons, including weaker recruitment during ipsilateral orienting toward a conspecific. In children with autism spectrum disorder, reduced functional connectivity between the SC and VTA is associated with the severity of social-orienting deficits^45^. Social investigation in rodents frequently involves close facial alignment, nose-to-nose contact, and strong protraction of multiple whiskers toward a conspecific^46–48^. The approximately symmetric whisker protraction evoked in our experiments could therefore represent a motor component that is also useful during social facial investigation. However, because our experiments did not include social stimuli, whether this response participates in social orienting or social motivation remains an open question.

### SC stimulation recruits temporally organized movement sequences

The temporal organization of the response argues against a unitary motor impulse delivered simultaneously to all effectors. Nose and whisker movements generally preceded eye movements, but their relative order varied across trials (**Figure 3A-C**). Three sequences accounted for most responses, indicating a constrained repertoire rather than one invariant chain. This combination of stereotypy and variability resembles natural orofacial behavior. Whisker positioning, eye movements, attempted head movements, and other facial movements are coupled during spontaneous behavior, but their timing and degree of coordination vary across actions and behavioral conditions^26–28^. Our results show that a similar pattern of coordination and variability can be initiated by perturbing the SC and remains apparent under head fixation, when the head and body components of orienting cannot be expressed.

This organization is also consistent with recent analyses of natural gaze shifts in freely moving mice. Natural gaze shifts comprise several reproducible eye-head coordination motifs that differ in the relative timing and contributions of the eyes and head^49^. Recordings in freely moving mice likewise show that SC activity is strongly modulated around gaze shifts and other movements^25^. Our stimulation experiments reveal a similar organizational principle from the opposite direction: beginning with a defined SC perturbation, the resulting orienting response was assembled from several effectors whose timing varied among a limited set of motifs. Together, the natural and stimulation studies support a view of mouse orienting as a distributed behavioral response rather than an isolated eye or head movement.

### Activation of SC inhibitory neurons evoked contraversive movements

Unilateral activation of the SC is classically associated with movements toward contralateral space^8,9,12,19,39,50^, whereas suppression of SC activity generally biases movements toward the ipsilateral side^50,51^. The consequences of selectively activating SC GABAergic neurons are less straightforward. Such stimulation can bias spatial choices or movements contraversively, as observed by Essig et al.^35^ and in the present study, but can also produce ipsiversive movements under other experimental conditions^52^. These differing effects likely reflect the heterogeneity of SC GABAergic neurons, which include local, commissural, and other long-range projecting populations^40^, as well as differences in the neuronal populations, collicular locations, and behavioral contexts examined.

Essig et al.^35^ found that activating SC GABAergic neurons biased choices contraversively without producing the local suppression expected from a simple inhibitory model. Their computational analysis showed that this result could arise when inhibition between the two colliculi dominates over inhibition within the stimulated SC. Our findings are compatible with such a network-level mechanism. Activating GABAergic neurons in one SC produced contraversive horizontal eye and nose movements and heterogeneous responses in the opposite SC. Moreover, delayed contralateral excitation, rather than the initial short-latency response, was associated with whether stimulation recruited an overt eye or nose movement. These observations suggest that the behavioral effect cannot be understood solely as local inhibition within the stimulated SC. Instead, stimulation appears to initiate an evolving bilateral network response whose later activity is more closely related to movement recruitment. Because we did not record from the stimulated SC, however, we cannot determine whether the stimulation produced local inhibition, disinhibition, or a mixture of local and long-range effects.

Finally, these findings clarify what activation of SC GABAergic neurons could represent and what it may not represent. Activating a genetically broad inhibitory population is unlikely to reproduce the physiological activity pattern that normally precedes orienting. The manipulation may recruit local inhibition, long-range GABAergic projections, disinhibition, antidromic activity, and secondary network responses. The evoked behavior should be interpreted as the consequence of perturbing an SC circuit, not as evidence that GABAergic neurons encode or generate the complete response. Nevertheless, the structure of the evoked output is informative. A relatively focal and temporally defined perturbation was sufficient to recruit multiple effectors, demonstrating that the wider sensorimotor network can transform local SC activation into a coordinated orienting action.

## Acknowledgements

We would like to thank the Charite workshop in particular Jan-Erik Ode, Alexander Schill and Daniel Deblitz for machine and electronics work.

## Author Contributions

HS, TL, MH, JS performed research. ML, RS, JK designed research. HS, MH, JS, KLT, KL, JK analyzed data; HS, RS, JK wrote the first draft of the paper; HS, JS, RS, JK edited the paper.

## Data availability

Raw data or code used are available upon requests.

## Supplementary materials

**Supplementary figure 1.**
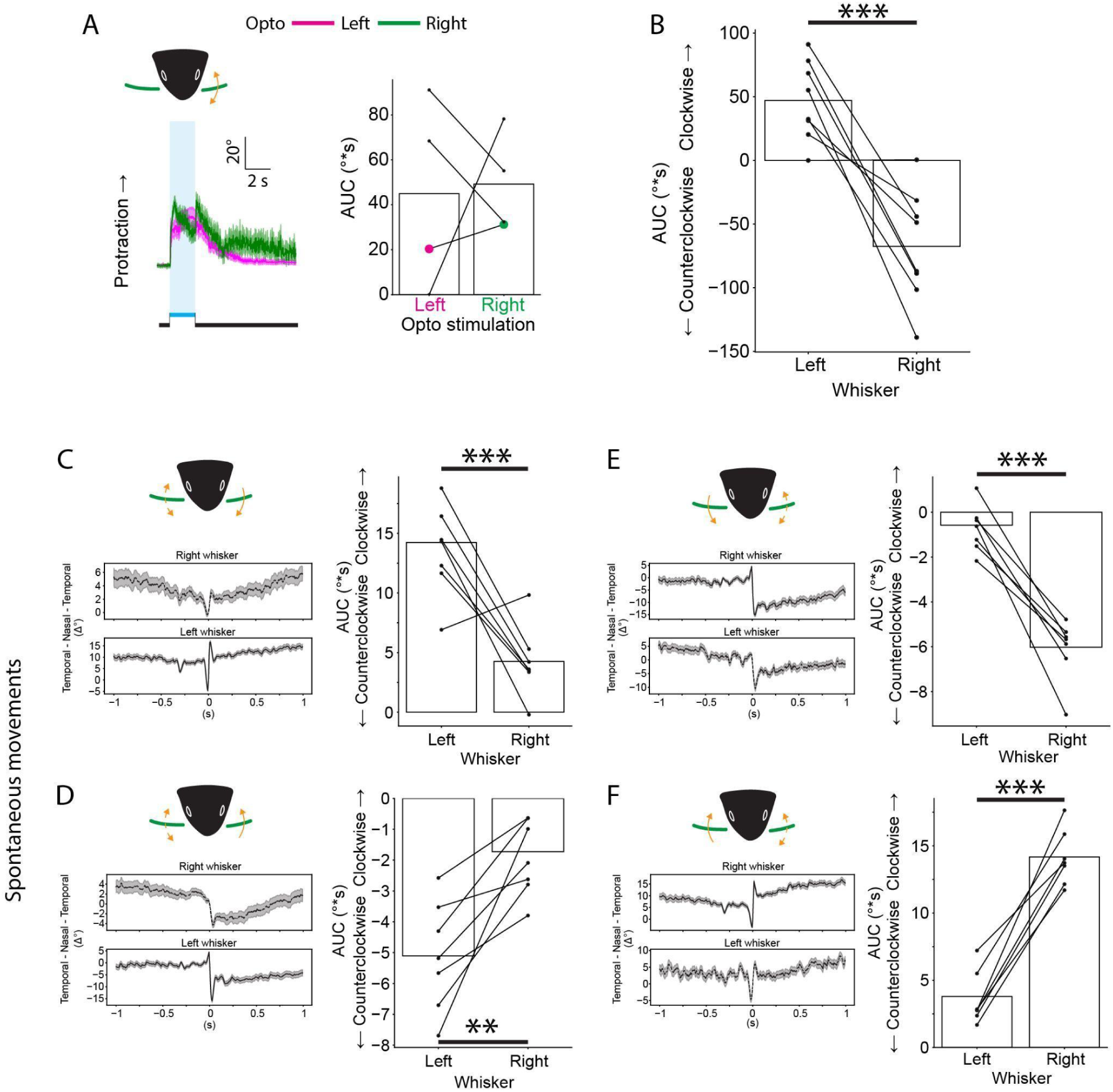
Asymmetric whisker movement evoked by optogenetic stimulation of the SC and symmetric spontaneous whisker movement of head-fixed mice on the moving platform. **(A)** Whisker movements following left and right SC stimulation, measured from the right whisker field. Left, mean whisker angle; right, AUC for individual animals. Whisker protraction showed no significant dependence on stimulation side. Traces show mean ± SEM across animals. Blue shading and bars indicate the optogenetic stimulation period. Dots and connecting lines represent individual animals; magenta and green symbols indicate group means for left and right SC stimulation, respectively. **(B)** AUC of evoked movements of right and left whiskers. Note that positive and negative values represent clockwise and counterclockwise, respectively. **(C-F)** Spontaneous whisker movement. Left, mean traces ± SEM; right, AUC for individual animals. Traces were aligned to left whisker protraction (C), left whisker retraction (D), right whisker protraction (E), right whisker retraction (F). Values were compared using paired t.test. **, p<0.01; ***, p<0.001.

**Supplementary figure 2.**
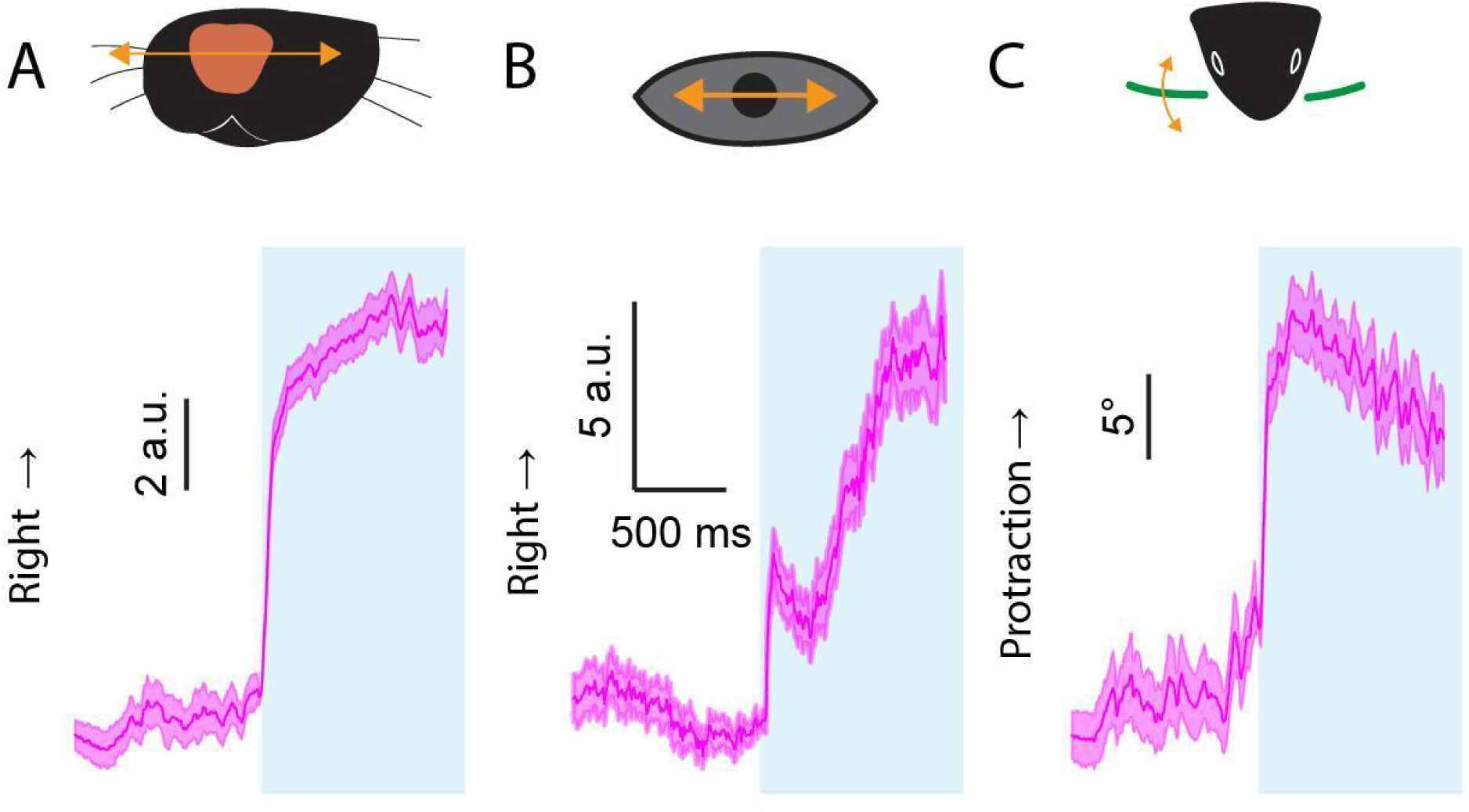
Optogenetic stimulation evoked orofacial movement of neuropixels recording experiments. **(A-C)** Mean traces ± SEM of optogenetic stimulation (left hemisphere) evoked horizontal eye movement (A), horizontal nose movement (B), whisker protraction (C) in a representative session. Blue shades represent optogenetic stimulation.

**Supplementary figure 3.**
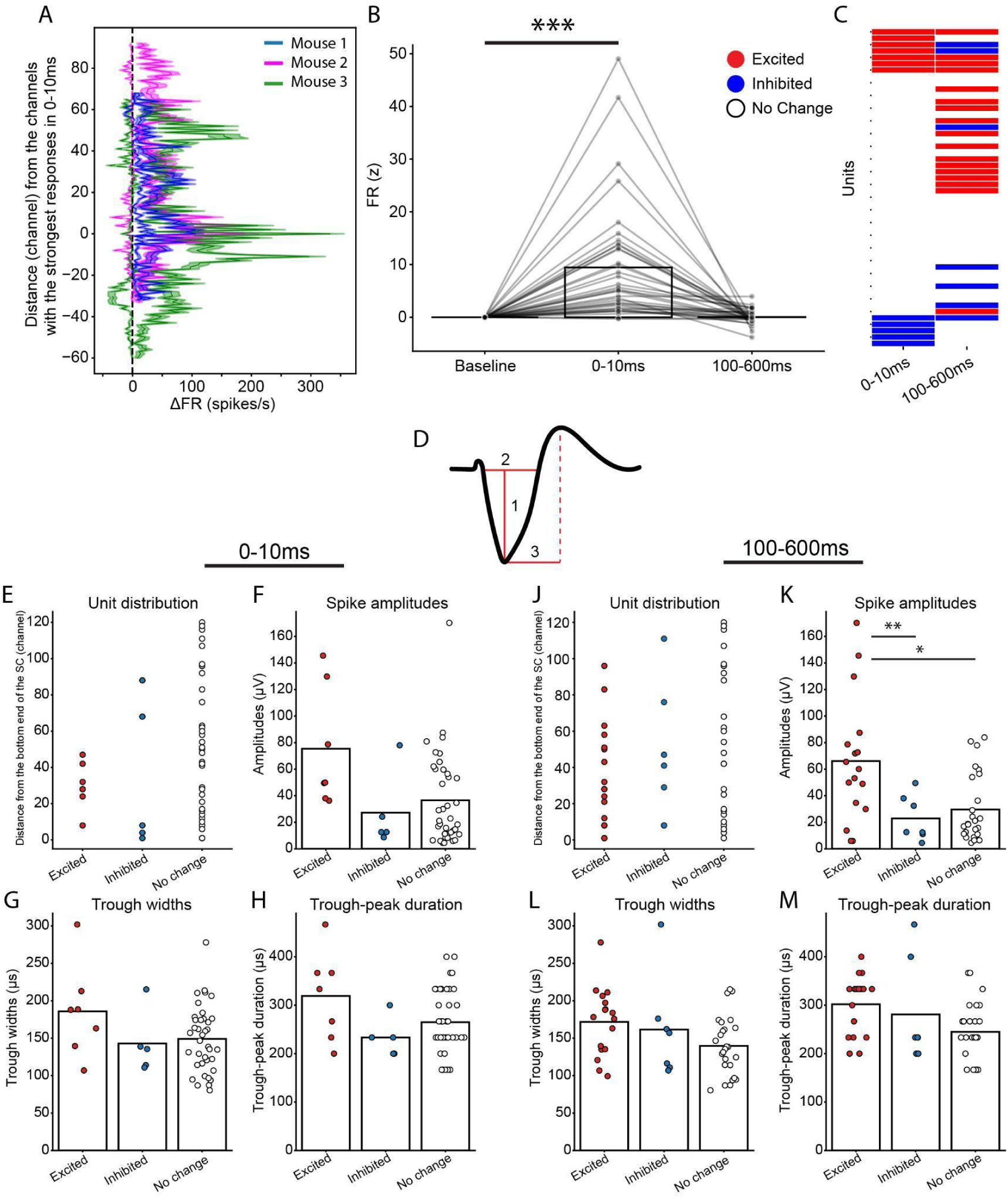
MUA and spike properties of single units. **(A)** Distribution of optogenetic stimulation evoked MUA responses. Solid and dotted lines show mean fast (0-10 ms) and late (100-600 ms) responses ± SEM aligned to the channel responded to stimulation the most strongly in a 0-10 ms time window. Colors represent individual animals. **(B)** Quantified MUA firing rates of every 10 channels. Comparison was performed with Friedman test followed by Wilcoxon signed-rank test with Holm-Bonferroni correction. **(C)** Statistical significance of single unit responses to optogenetic stimulation. Red, excited; blue, inhibited; white, significantly not changed. Significance is when p<0.05. **(D)** Schematic action potential showing spike properties that we quantified. 1, spike amplitudes; 2, trough widths; 3, trough-peak duration. **(E-M)** Spike properties of excited, inhibited, not changed units in a 0-10 ms window (E-H) and in a 100-600 ms window (J-M). Properties include unit distribution (E,J), spike amplitudes (F,K), trough widths (G,L), and tough-peak duration (H,M). Comparison was performed with LMM accounting for animal identities followed by MWU with Holm-Boferroni correction. *p < 0.05, **p < 0.01, ***p < 0.001.

**Supplementary video 1. Representative video of right and left eyes in a right SC optogenetic stimulation trial.** Left, right eye; right, left eye. Right SC stimulation evoked leftward eye movement (direction to right in the view) in both eyes. The same trial as Supplementary video 2 and 3.

**Supplementary video 2. Representative video of front view in a right SC optogenetic stimulation trial.** Right SC stimulation evoked leftward nose movement (direction to right in the view). The same trial as Supplementary video 1 and 3.

**Supplementary video 3. Representative video of top view in a right SC optogenetic stimulation trial.** Whiskers were painted with UV-sensitive dye, green in the video. Blue signal is optogenetic LED stimulation at 470 nm. Right SC stimulation evoked whisker protraction (direction to left in the view). The same trial as Supplementary video 1 and 2.

